# Morphofunctional Characterization of Colonic Organoids from Muc2^−^/^−^ Mice

**DOI:** 10.64898/2026.10.21.760621

**Authors:** J. V. Popova, S. S. Medvedeva, A.V. Zybkina, E. N. Kozhevnikova

## Abstract

Inflammatory bowel diseases (IBD) remain a serious challenge in gastroenterology, and disruption of the mucus barrier integrity plays a key role in their pathogenesis. The Muc2 protein is the main structural component of the protective mucus layer in the colon, and mice with a knockout of the *Muc2* gene (*Muc2*^−^/^−^) serve as a classic model for studying IBD. In the present work, three-dimensional cultured organoids derived from the colonic crypts of *Muc2*^−^/^−^ mice and their wild-type littermates (*Muc2*^+^/^+^) were used for a comparative analysis of their morphofunctional characteristics. Using morphometric analysis and flow cytometry to identify the Lgr5^+^ stem cell population, we demonstrated that organoids obtained from *Muc2*^−^/^−^ animals are characterized by a smaller size, absence of crypt-like structures, increased proliferative activity, and a reduced proportion of stem cells compared to organoids derived from wild-type littermates. The obtained data indicate that Muc2 deficiency and the associated intestinal inflammation affect epithelial architecture and stem cell balance outside the inflammatory environment *in vitro*. Thus, 3D colonic organoids derived from *Muc2* knockout mice represent a promising tool for studying IBD and testing novel therapeutic strategies.

## INTRODUCTION

Inflammatory bowel diseases (IBD), including ulcerative colitis and Crohn’s disease, are chronic, relapsing inflammatory disorders of the gastrointestinal tract whose prevalence is steadily increasing in industrialized countries and reaches 0.5–1% of the population [1, 2]. The etiology of IBD remains multifactorial, involving genetic predisposition, dysregulation of innate and adaptive immunity, alterations in the gut microbiota, and disruption of epithelial barrier integrity. One of the key elements of IBD pathogenesis is considered to be increased intestinal barrier permeability, which leads to enhanced penetration of bacterial antigens into the underlying lamina propria and to the initiation of chronic immune-mediated inflammation [3, 4].

A critical role in maintaining the barrier function of the colon is played by the mucus layer covering the epithelium. The main structural component of this layer is the high-molecular-weight glycoprotein Muc2, which is secreted by goblet cells and forms a gel-like polymeric network owing to its ability to oligomerize through cysteine-rich domains [5, 6]. The Muc2 protein forms two structurally and functionally distinct layers: an inner layer, firmly attached to the epithelium and essentially devoid of bacteria, which serves as the main physical barrier, and an outer, loose layer colonized by commensal microbiota that metabolize mucin oligosaccharides. Thus, the Muc2 protein acts not only as a passive physical barrier but also as an active regulator of the interaction between the microbiota and the host immune system [5, 6].

One of the experimental models for studying the role of the Muc2 protein and IBD in general is the *Muc2* knockout mouse (*Muc2*^−^/^−^) [7]. These animals spontaneously develop colitis that resembles human ulcerative colitis in its clinical and histological features, with characteristic manifestations: weight loss, diarrhea, rectal bleeding, inflammatory cell infiltration, crypt abscess formation, and superficial erosions. In *Muc2*^−^/^−^ mice the mucus layer is absent, which results in direct contact of bacteria with epithelial cells, increased production of proinflammatory cytokines (TNF-α, IL-6, IL-1β), and activation of signaling pathways associated with Toll-like receptors (TLRs) and NF-κB [8]. In addition, pronounced hyperproliferation is observed in the crypt epithelium of these animals, which is thought to represent a compensatory response to chronic tissue damage [7, 9]. However, when interpreting data obtained *in vivo*, uncertainty always remains: which changes are a direct consequence of the absence of the Muc2 protein in the epithelium itself, and which are secondary, driven by chronic inflammation and the altered composition of the gut microbiota. To address this question, earlier studies were performed on germ-free *Muc2*^−^/^−^ and *Muc2*^+^/^+^ mice, which showed that epithelial hyperproliferation persists in the absence of bacteria and is also reproduced in organoid cultures. This indicates that the altered phenotype of intestinal cells does not depend on external stimuli, although the influence of the surrounding tissues still remains strong [9].

A breakthrough in the study of intestinal cell biology was the development of 3D organoid cultures. The technology, first described by Hans Clevers’ group, allows three-dimensional structures to be obtained from single Lgr5^+^ stem cells or from isolated crypts, which, in the presence of growth factors (EGF, Noggin, R-spondin-1) and extracellular matrix (Matrigel), self-organize into structures resembling the intestinal epithelium in architecture [10–12]. Organoids maintain the hierarchy of cell types: stem cells (Lgr5^+^) reside in the region corresponding to the bottom of the crypt, whereas transit-amplifying (TA) cells and differentiated enterocytes, goblet cells, and enteroendocrine cells are located in the apical part. The main advantage of organoids is the possibility to grow such three-dimensional “mini-organs” under controlled conditions, fully isolated from bacteria, immune cells, and blood factors. This allows one to see in its pure form how genetic defects change the structure and functioning of the intestinal epithelial tissue [10, 13].

In the context of studying the role of the Muc2 protein, as well as in IBD research in general, the organoid model makes it possible to investigate the effects of chronic inflammation in isolation. The use of these three-dimensional cultures helps to understand which alterations persist in the intestinal tissue after prolonged Muc2 deficiency, outside the systemic influence of the organism. Although pilot studies revealed hyperproliferation of organoids derived from *Muc2*^−^/^−^ mice [9] and instability of their phenotype during long-term culture [14], a comprehensive analysis of the spatial organization of the actin cytoskeleton and FACS analysis of the cell composition has not previously been performed. Understanding how mucin deficiency affects the organization of the actin cytoskeleton, which is required for maintaining apico-basal polarity, tight junctions, and the mechanical integrity of the epithelium, is of key importance [15, 16]. Moreover, whereas tight junction defects and disruption of apical F-actin under mucin deficiency have already been described at the level of whole tissue *in vivo* [17], the pattern of structural changes in actin in 3D organoids remains unexplored. In addition, the changes in the proliferative balance and in the composition of stem and transit-amplifying cell populations under conditions of culturing mucin-deficient epithelium require clarification, especially in light of the data indicating that signaling cascades regulating the stem cell niche (including Wnt) are involved in the control of epithelial proliferation [18].

Since proliferative activity, barrier function, and the organization of the actin cytoskeleton under mucin-2 deficiency are difficult to assess correctly in monolayer cultures consisting of a single epithelial cell type, we used intestinal organoids, which reproduce the three-dimensional architecture of the epithelium and contain the full spectrum of cell populations. We hypothesized that if organoids derived from the crypts of *Muc2*^−^/^−^ mice (a model of chronic colitis) retain increased proliferative activity outside the organism, then this effect should be associated with changes in the number or activity of stem cells (Lgr5^+^). To test this hypothesis, a systematic comparative analysis of the morphofunctional properties of organoids between the *Muc2*^−^/^−^ and *Muc2*^+^/^+^ groups was performed, with assessment of growth, epithelial barrier permeability, and stem cell population composition using morphometry and flow cytometry (FACS).

## MATERIALS AND METHODS

### Laboratory animals and housing conditions

Animal experiments were carried out at the Institute of Molecular and Cellular Biology SB RAS and the Siberian State University of Engineering and Biotechnology (formerly NSAU). All research procedures involving laboratory animals complied with the ethical standards approved by the legal acts of the Russian Federation and with the principles of the Basel Declaration. The experimental protocols were approved by the IMCB SB RAS Ethics Committee (protocol No. 02/21 of 04/08/2021).

The study was performed on animals homozygous for the *Muc2* gene knockout (*Muc2*^−^/^−^, n = 3). Wild-type littermates (*Muc2*^+^/^+^, n = 3) were used as controls. The animals were housed at the testing center of the Siberian State University of Engineering and Biotechnology (formerly NSAU) in same-sex groups of 5–8 animals in open cages. The cages were provided with bedding made of dust-free birch shavings (Fraction No. 3, IP Filonich, Russia) and cardboard shelters (cardboard tubes). The animals had ad libitum access to food (C-19 feed, Delta Feeds, BioPro, Russia) and filtered water. The animal housing room maintained a 12:12 light cycle (lights on at 24:00, lights off at 12:00), a temperature of 22–24°C, and a humidity of 30–60%.

### Isolation of crypts and cultivation of colonic organoids

Crypt isolation and organoid cultivation were performed according to a modified protocol of Sato et al. [11, 12]. The descending colon was excised, washed with cold PBS without Ca^2+^/Mg^2+^ (dPBS; cat. no. CP060-10, PanEco, Russia) using a catheter, cut longitudinally, and minced into ∼2 mm fragments. The tissue was washed twice in 3 ml of dPBS until the supernatant became clear. To isolate crypts, the tissue was incubated in 5 ml of chelation buffer consisting of dPBS, 2 mM EDTA (cat. no. 1108KG001, Macklin), and 43.4 mM sucrose for 30 min at 4°C. The buffer was removed, 3–5 ml of fresh buffer was added, and the suspension was gently shaken 5–6 times. The supernatant (crypts) was collected, and the procedure was repeated 4–5 times. Fractions containing single cells were not used. The combined fractions were centrifuged (200 g, 5 min, 4°C), and the pellet was resuspended in 1 ml of wash medium consisting of DMEM/F12 (cat. no. bn-3A9A, BioinnLabs, Russia) and 10% FBS (cat. no. RM10432, HiMedia); centrifugation was repeated. The crypt pellet was resuspended in 100 µl of wash medium.

To assess the number and viability of crypts, 9 µl of suspension was mixed with 1 µl of 0.4% trypan blue (cat. no. 72-57-1.25, Suvchem), and unstained crypts were counted. The volume of suspension containing 50–150 crypts per well of a 48-well plate (cat. no. 92048, TPP) was calculated. The selected volume was centrifuged (200 g, 5 min, 4°C), and the pellet was resuspended in DMEM/F12 + 10% FBS in a volume providing a final concentration of 50% Cultrex matrix (cat. no. 3533-005-02, R&D Systems) in a final volume of 25 µl per well (12.5 µl cell suspension + 12.5 µl matrix). The mixture was gently homogenized with a wide-bore pipette tip (5–10 times), and 25 µl of suspension was placed in the center of each well of a plate pre-warmed to 37°C. After matrix polymerization (10–15 min), 250 µl of growth medium was added per well. The growth medium based on DMEM/F12 (cat. no. bn-3A9A, BioinnLabs, Russia) contained 50% L-WRN conditioned medium (a source of Wnt-3a, R-spondin-3, and Noggin) (cat. no. SCM105, Sigma), 10% FBS (cat. no. RM10432, HiMedia), 2% 50× B27 supplement / NeuroMax (cat. no. ФP-0305, PanEco, Russia), 1% 100× N2 supplement (cat. no. ФP-0205, PanEco, Russia), 500 µM N-acetyl-L-cysteine (cat. no. A9165, Sigma), 50 ng/ml EGF (cat. no. ФP-08000, PanEco, Russia), 10 µM ROCK inhibitor (Y-27632) (cat. no. 129830-38-2, Tocris Bioscience), and 1% antibiotic/antimycotic solution (ABM; cat. no. A5955, Sigma). The prepared medium was filtered through a sterile filter (0.22 µm, cat. no. FPE204025, Jet BIOFIL), stored at 4°C, and replaced every 48 h.

### Immunohistochemical analysis of colonic organoids

On day 7 of organoid culture, the matrix droplets were mechanically disrupted, the suspension was collected and fixed in 10% buffered formalin solution (cat. no. 3857.1000, LabElement, Russia) for 1 h at room temperature. Membrane permeabilization was performed for 1 h in buffer containing 0.3% Triton X-100 (cat. no. X-100-100ML, Sigma) and 0.5% BSA (cat. no. BSA.0100, Dia-M, Russia) in PBS. To visualize F-actin, samples were incubated with fluorescently labeled phalloidin (Phalloidin-TRITC, working concentration 1 µM; cat. no. P1951, Sigma) for 1 h in the dark. Cell nuclei were stained with DAPI (1:1000; cat. no. TC229-10MG, HiMedia). The stained organoids were gently mounted between a glass slide and a coverslip in 50% glycerol (cat. no. sc-29095, ChemCrus) in PBS and imaged using a Zeiss LSM 710 confocal laser scanning microscope (Carl Zeiss, Germany) with ZEN 2012 software.

### Functional epithelial barrier permeability assay

The epithelial barrier permeability *in vitro* was assessed according to a previously published protocol [14]. On day 7 of organoid culture, the matrix was mechanically disrupted, the organoids were collected and centrifuged (300 g, 5 min, 4°C). To the organoid suspension, 100 µl of HBSS (cat. no. CP020-10, PanEco, Russia) containing the low-molecular-weight fluorescent tracer FITC-dextran (4 kDa; cat. no. FD4-1G, Sigma-Aldrich) at a final concentration of 2 mg/ml was added. As a positive control, EDTA (cat. no. 1108KG001, Macklin) at a final concentration of 5 mM was added to the HBSS to induce disruption of calcium-dependent intercellular junctions. Organoids were imaged 5 min after the addition of FITC-dextran with or without EDTA on an inverted Zeiss LSM 710 confocal laser scanning microscope (Carl Zeiss, Germany) in brightfield and in the green fluorescence channel using ZEN 2012 software.

### Morphometric measurements and image analysis in ImageJ (Fiji)

Fluorescence intensity and morphometric parameters were quantified using Fiji software [19]. Organoid size and shape were assessed from brightfield micrographs obtained on day 7 of culture. For each object, the maximum Feret diameter (Feret max diameter, µm), the mean epithelial wall thickness (Cell wall width, µm), and the solidity coefficient (Solidity, a.u.), calculated as the ratio of the object’s area to the area of its convex hull, were determined, where the convex hull is the minimal convex shape completely enclosing the object.

To assess the state of the actin cytoskeleton, the mean fluorescence intensity of the TRITC signal was measured on confocal Z-stacks of 9.69 µm thickness in two zones: along the apical (apical F-actin) and lateral (lateral F-actin) domains of the plasma membrane of the cells.

The quantitative assessment of barrier permeability (Permeability, a.u.) was calculated as the ratio of the FITC fluorescence intensity in the organoid lumen to the mean of three background regions around the organoid. For each experimental group, at least 30 randomly selected organoids with an intact wall from three independent experiments were analyzed.

### Flow cytometry (FACS)

To analyze the cell composition, organoids on day 7 of culture were dissociated into a single-cell suspension. The culture plate with organoids was placed on ice for 2 min, then the growth medium was carefully removed from the wells without damaging the matrix droplets. 0.8 ml of ice-cold Gentle Cell Dissociation Reagent (cat. no. 07174, STEMCELL Technologies) was added to each well and pipetted vigorously 10–15 times with a P1000 tip to completely disrupt the matrix. The suspension from the wells was combined in a 15 ml tube and incubated strictly on ice for 15–20 min. The sample was then centrifuged (300 g, 5 min, 4°C), the supernatant removed, and the pellet washed with 5 ml of ice-cold dPBS and centrifuged again under the same conditions.

For enzymatic dissociation, the pellet was resuspended in 0.5 ml of TrypLE Express (cat. no. 12605-01, Gibco) pre-warmed to 37°C with the addition of 10 µM ROCK inhibitor (Y-27632) (cat. no. 129830-38-2, Tocris Bioscience) and incubated at 37°C for 15 min. Every 2–3 min, the tube was removed and the organoids were mechanically dissociated by pipetting (15– 20 times using a P1000 or P200 tip). Dissociation was monitored by phase contrast by taking a 10 µl aliquot. The enzymatic reaction was stopped by adding 2.5 ml of ice-cold FACS buffer consisting of dPBS with 2% BSA (cat. no. BSA.0100, Dia-M, Russia) and 10 µM Y-27632. The cell suspension was filtered through a cell strainer with a 20 µm pore size (cat. no. 43-50020-01, PluriSelect), pre-moistened with FACS buffer, to remove aggregates and debris. The filtrate was centrifuged (450 g, 7 min, 4°C), and the supernatant removed. The pellet was resuspended in 500 µl of cold FACS buffer. The cell yield was approximately 50,000–100,000 cells per organoid well depending on density and genotype.

The resulting suspension was divided into two parts: the test sample and the negative control. Staining was performed on ice for 30 min in the dark. To the test sample, 2 µl of anti-mouse Lgr5 antibodies conjugated with APC (clone DA04-10E8-9, cat. no. 130-111-390, Miltenyi Biotec) and 2 µl of anti-mouse EpCAM antibodies conjugated with FITC (clone CAA7-9G8, cat. no. 130-123-674, Miltenyi Biotec) were added. To assess viability, 0.5 µl of DAPI (1 mg/ml, cat. no. TC229-10MG, HiMedia) was added to all samples. The negative control was stained with DAPI only for cytometer setup and determination of the viability threshold. After incubation, the cells were washed by centrifugation with buffer replacement to remove unbound antibodies. Analysis was performed on a NovoCyte flow cytometer (Agilent). Viable cells were identified as DAPI-negative, with epithelial (EpCAM^+^) and stem (Lgr5^+^) cells distinguished among them. The gating strategy included selection of the target cell population by FSC-H/SSC-H parameters (Cells), subsequent doublet exclusion in the SSC-H/SSC-A gate (Single cells), and final calculation of the percentage of stem cells in the Lgr5^+^EpCAM^+^ quadrant among viable epithelial cells (Lgr5^+^DAPI^−^). The results were expressed as the percentage of Lgr5^+^EpCAM^+^ cells of the total number of viable cells. At least 10,000 events were recorded per sample. The data were analyzed using FlowJo v10.8 software (BD).

### Statistical analysis

Statistical processing of the quantitative data was performed using R programming language packages (version 4.4.1), and the graphs were visualized using the ggplot2 library. The distribution of the obtained data was assessed using the Kolmogorov–Smirnov test. For comparison of two independent groups (*Muc2*^+^/^+^ and *Muc2*^−^/^−^), the unpaired two-sample Student’s *t*-test was applied for normally distributed data, Welch’s *t*-test for two independent samples with small group sizes, and the nonparametric Mann–Whitney *U*-test when the data distribution deviated from normal. Differences were considered significant at *p* < 0.05. The data in the graphs are presented as boxplots combined with dot plots of individual measurements. Statistically significant differences in the graphs are marked with asterisks: * ‒ *p* < 0.05, ** ‒ *p* < 0.01, *** ‒ *p* < 0.001.

## RESULTS

### Comparative morphometric analysis of the growth dynamics of colonic organoids from Muc2^+^/^+^ and Muc2^−^/^−^ mice

At the initial stage of the work, a comparative assessment of the growth and morphological features of 3D colonic organoid cultures derived from *Muc2*^+^/^+^ mice and from animals with a knockout of the gene encoding mucin-2, *Muc2*^−^/^−^, was performed. Observation of the development of the cultures in brightfield over 7 days of cultivation revealed marked differences in their spatial organization (Fig. 1a). Organoids derived from littermates without the mutation (*Muc2*^+^/^+^) demonstrated a classic pattern of development: by day 7 of incubation they branched, forming numerous crypt-like outgrowths. In contrast, organoids derived from the epithelium of *Muc2*^−^/^−^ mice remained predominantly spherical throughout the entire observation period and showed virtually no branching (Fig. 1a). Moreover, most *Muc2*^−^/^−^ organoids contained dense material in their lumen, representing an accumulation of cellular debris composed of sloughed epithelial cells.

**Fig. 1.**
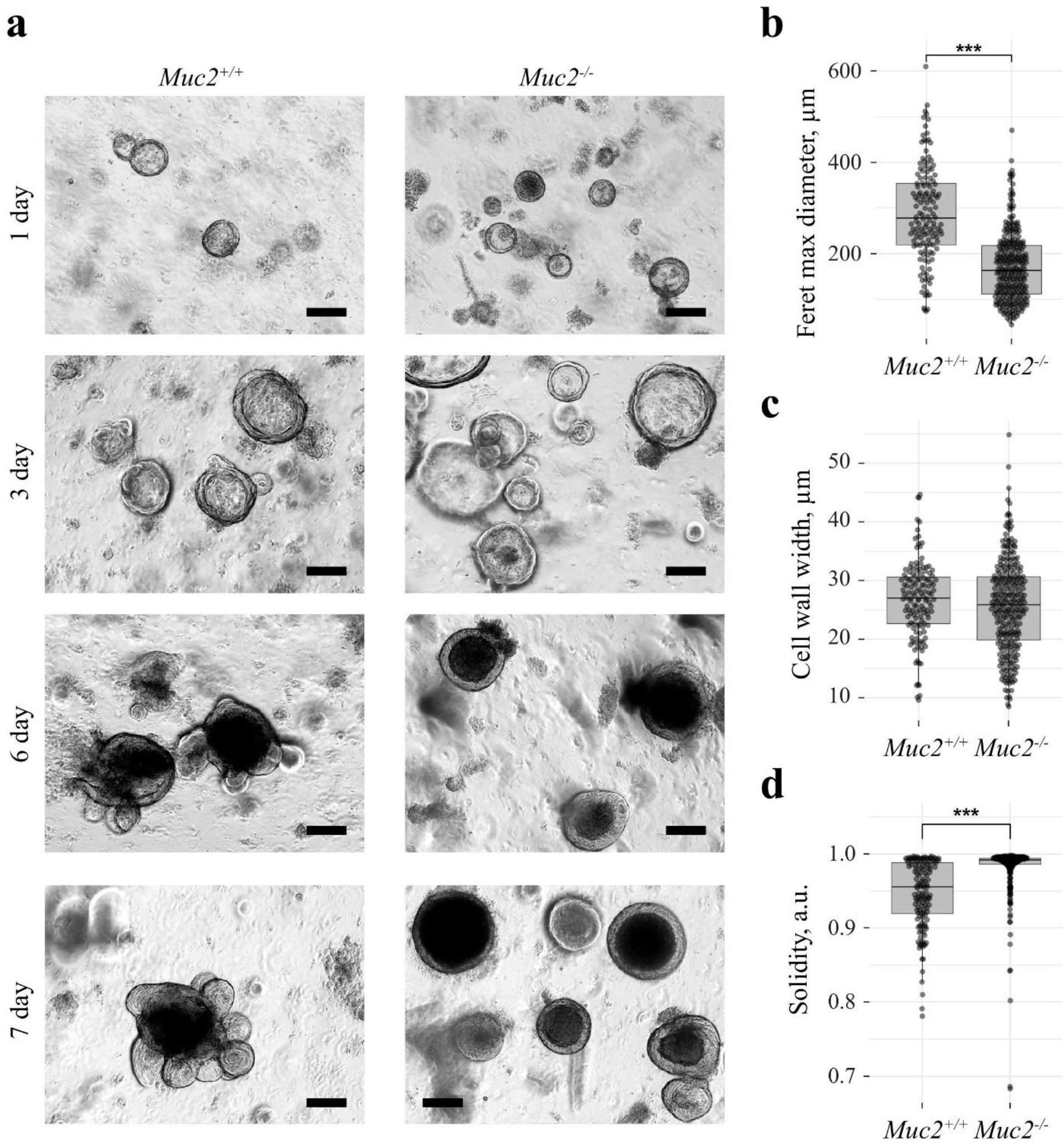
Morphometric characteristics of colonic organoids from *Muc2*^+^/^+^ and *Muc2*^−^/^−^ mice over 7 days of culture. (a) Brightfield micrographs of three-dimensional cultures on days 1, 3, 6, and 7 of cultivation; scale bar ‒ 100 µm. (b) Distribution of the maximum Feret diameter (Feret max diameter, µm). (c) Organoid wall thickness (Cell wall width, µm). (d) Contour roundness index (Solidity, a.u.). Data are presented as boxplots overlaid with individual measurement points; *** ‒ *p* < 0.001. Student’s *t*-test was used for (b, c); Mann–Whitney *U*-test was used for (d).

To verify the observed changes, a morphometric analysis of the cultures was performed on day 7 of the experiment. Measurement of the maximum Feret diameter showed a statistically significant (*p* < 0.001) decrease in this parameter in *Muc2*^−^/^−^ organoids compared to the control (Fig. 1b): the mean diameter values were about 150–170 µm for cultures derived from animals with the mutation versus 250–280 µm for cultures derived from their mutation-free *Muc2*^+^/^+^ littermates. At the same time, assessment of the organoid wall thickness (Cell wall width) revealed no significant differences between the groups studied (Fig. 1c), remaining in the range of 25–30 µm in both groups, which indicates a largely preserved columnar shape of the cells.

Important quantitative confirmation of the phenotypic differences came from the data on the Solidity parameter (roundness factor), which reflects the degree of contour indentation of an object. Organoids derived from *Muc2*^−^/^−^ mice were characterized by significantly higher Solidity values (*p* < 0.001), clustered in the range of 0.95–1.00 (Fig. 1d). At the same time, organoids derived from wild-type animals showed a wide distribution of this parameter with a marked shift toward lower values (down to 0.8–0.9 a.u.). This is due to the active formation of peripheral outgrowths and the increasing complexity of their three-dimensional structure. Thus, the epithelium derived from *Muc2* knockout mice forms stable, symmetric spheroids of smaller diameter with suppressed morphogenetic budding in organoid culture.

### Assessment of epithelial barrier permeability and of the proportion of stem cells in organoids derived from Muc2^+^/^+^ and Muc2^−^/^−^ mice

To assess the integrity of intercellular junctions *in vitro*, a functional permeability test was performed using the fluorescent tracer FITC-dextran 4 kDa (Fig. 2a). EDTA, which transiently disrupts calcium-dependent tight junctions, was used as a positive permeability control. Under control conditions (CTR), organoids derived from both groups of animals were impermeable to the dye. This indicates that the presence of the Muc2 protein is not in itself required for the formation of the epithelial barrier. At the same time, the increased intestinal barrier permeability characteristic of *Muc2*^−^/^−^ animals is not preserved when the epithelium is cultured *in vitro*, pointing to the key role of inflammatory environmental factors in barrier dysfunction.

**Fig. 2.**
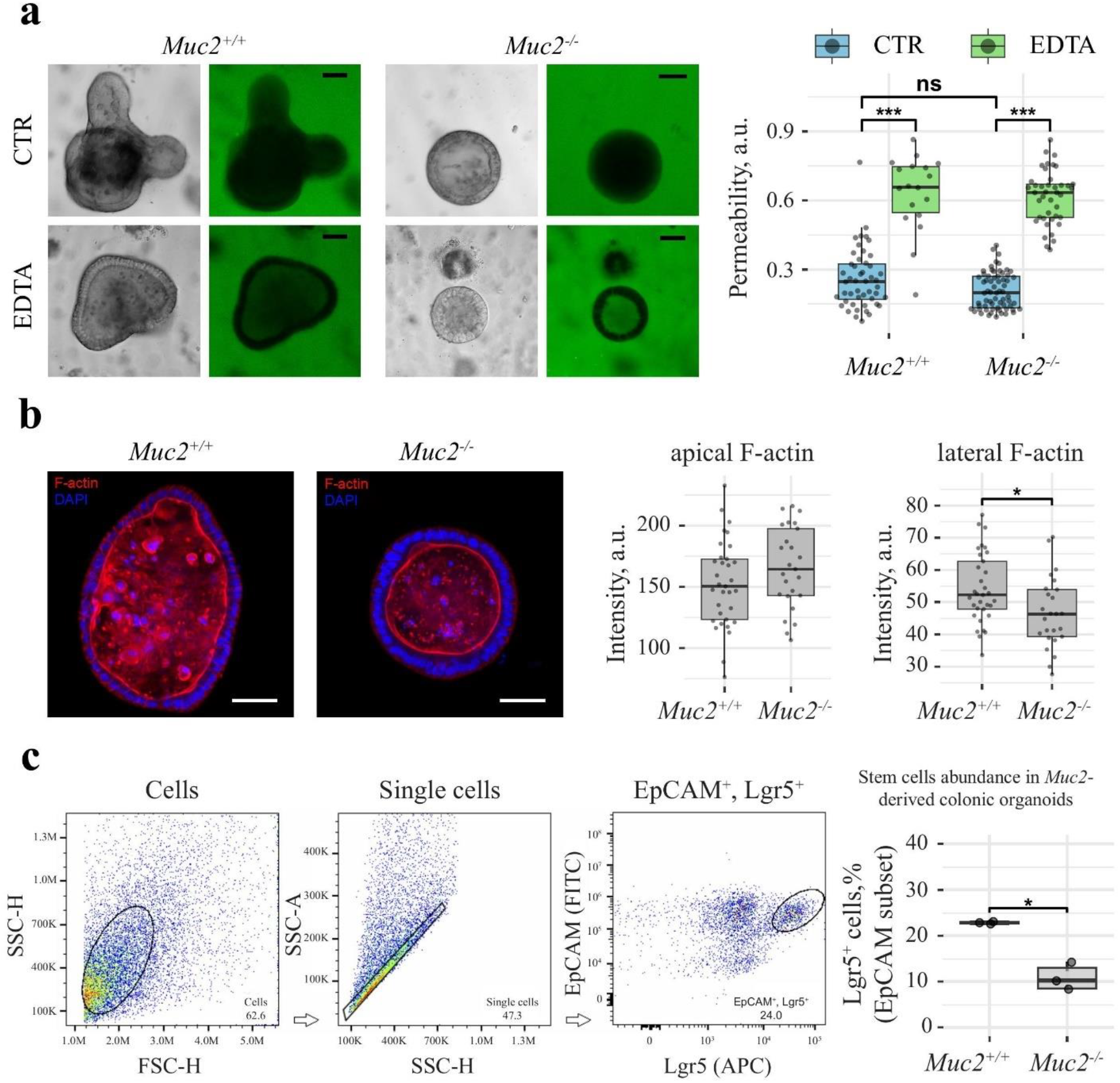
Barrier properties, F-actin organization, and FACS analysis of stem cells in colonic organoids from *Muc2*^+^/^+^ and *Muc2*^−^/^−^ mice. (a) Brightfield and fluorescence micrographs showing the epithelial barrier permeability assay under control conditions (CTR) and after EDTA treatment; scale bar ‒ 50 µm. Right: quantification graph of barrier permeability (Permeability, a.u.). (b) Confocal images of organoids stained for F-actin (red) and nuclei with DAPI (blue); scale bar ‒ 50 µm. Right: fluorescence intensity graphs of apical and lateral F-actin. (c) Gating strategy for single viable epithelial cells and distribution of the stem cell population based on EpCAM (FITC) and Lgr5 (APC) markers. Right: graph showing the proportion of Lgr5^+^ cells, % within the EpCAM^+^ population. * ‒ *p* < 0.05, *** ‒ *p* < 0.001, ns ‒ non-significant; Welch’s *t*-test.

Next, the spatial organization of the actin cytoskeleton (F-actin) was studied by confocal microscopy (Fig. 2b). Quantitative analysis of the fluorescent signal distribution showed that the intensity of apical actin (apical F-actin), which forms the brush border, remained stable and did not differ significantly between the *Muc2*^+^/^+^ and *Muc2*^−^/^−^ groups. However, in the lateral domains of the plasma membrane (lateral F-actin), a statistically significant (*p* < 0.05) decrease in fluorescence intensity was found in the epithelium of the *Muc2*^−^/^−^ organoid group. Such changes are characteristic of animals of this genotype and are partially preserved when the epithelium is cultured *in vitro* (Fig. 2b).

To assess the proportion of stem cells in the organoids of both groups, flow cytometry (FACS) of stem cells was performed using Lgr5^+^EpCAM^+^ markers (Fig. 2c). In organoids derived from *Muc2* knockout mice, a twofold and statistically significant (*p* < 0.05) decrease in the intestinal stem cell pool was found: the proportion of Lgr5^+^ cells among EpCAM^+^ epithelial cells dropped from ∼23% in organoids derived from wild-type mice to ∼10% in organoids derived from *Muc2*^−^/^−^ animals. Thus, the increased proliferation of cells outside the stem niche, characteristic of the *Muc2*^−^/^−^ epithelium *in vivo*, is also preserved during organoid culture.

## DISCUSSION

The data obtained in the present study reveal new fundamental aspects of the understanding of the role of mucin-2 in the regulation of intestinal epithelial biology. The use of three-dimensional (3D) organoid cultures made it possible to study the features of the *Muc2* knockout epithelium while completely excluding the concomitant systemic factors present *in vivo*, such as tissue infiltration by immune cells, exposure to gut microbiota metabolites, and the systemic pool of proinflammatory cytokines. This approach made it possible to distinguish the intrinsic rearrangements of the epithelium under conditions of mucus layer damage from the secondary effects induced by chronic inflammation.

One of the key morphological phenotypes was the pronounced alteration of the spatial structure of the organoids from animals with chronic intestinal inflammation. Organoids derived from *Muc2*^−^/^−^ mice demonstrated a significant decrease in the maximum Feret diameter along with a sharp increase in the Solidity parameter (Fig. 1b, d). This quantitatively confirms that organoids from *Muc2*^−^/^−^ mice retain a spherical shape throughout all 7 days of culture and have a reduced capacity to form crypt-like outgrowths. The classical studies of H. Clevers’ group showed that the branching of intestinal organoids *in vitro* is determined by the structure of the stem cell niche and the uneven distribution of Wnt/β-catenin and Notch signaling pathway activity [10–12]. The stable maintenance of a spherical shape by organoids usually indicates either a disruption of local morphogen gradients or a change in proliferative status and loss of the pool of true stem cells.

This conclusion finds direct quantitative confirmation in the results of our study, which revealed a pronounced twofold decrease in the proportion of Lgr5^+^EpCAM^+^ stem cells in organoids derived from *Muc2*^−^/^−^ animals compared to the control group (Fig. 2c). Previously, *in vivo* studies in *Muc2* knockout mice described pronounced cell proliferation in the middle and upper parts of the colonic crypts [20, 21]. The fact that the Lgr5^+^ stem cell pool is reduced under culture conditions indicates that the *in vivo* crypt hyperplasia is most likely not determined solely by the properties of the epithelium itself. The increased proliferation in *Muc2*^−^/^−^ mice may be compensatory in nature, in response to chronic damage to the epithelium unprotected by the mucus layer [8, 20]. However, this phenotype persists in germ-free animals and in organoids, indicating its independence from microbial ligands (via TLR- and NF-κB-dependent pathways) and from the stromal microenvironment (myofibroblasts, endothelial cells) [9]. In the *in vitro* epithelial organoid system, devoid of bacteria and immune cells, *Muc2* knockout is accompanied by a reduction of the Lgr5^+^ cell pool. This phenomenon may be underlain by a disruption of the spatial distribution of growth factors supporting the stem cell niche. The importance of proper regulation of niche factors for the proliferative status of the crypts is well known [18]; however, the role of the extracellular mucin matrix in this process remains unclear. We propose that mucin-2, secreted by epithelial cells, participates in the formation of the crypt microenvironment required for retaining and presenting paracrine growth factors, in particular Wnt ligands, on the cell surface. In the absence of the mucin matrix, the availability of these factors to their receptors (Frizzled, Lgr5) may decrease, leading to weakening of the Wnt/β-catenin cascade, disruption of the symmetry of the epithelial sheet, and suppression of budding. This is consistent with the data on the gradual destabilization of the epithelial phenotype during long-term culture of *Muc2*-deficient organoids [14]. Apparently, the chronic inflammation experienced *in vivo* leaves a trace in the subsequent proliferation and differentiation of epithelial cells, which manifests itself as exhaustion of the stem cell pool even under culture conditions.

Another key finding of the work was the discovery of a selective defect of the actin cytoskeleton. Quantification of the fluorescent signal showed that, with unchanged levels of apical F-actin forming the brush border, the intensity of lateral F-actin in organoids from *Muc2*^−^/^−^ mice was significantly reduced (Fig. 2b). The lateral, juxtamembrane actin framework plays a critical role in stabilizing the complexes of intercellular junctions ‒ adherens junctions (mediated by E-cadherin) and tight junctions (based on ZO-1 family proteins and claudins) [15, 16]. The cytoskeleton provides mechanical coupling of the cells and transmission of biomechanical signals within the epithelial sheet.

Previously, at the level of whole tissue in mice *in vivo*, profound ultrastructural disruption of tight junctions accompanied by mitochondrial dysfunction has been described under Muc2 deficiency [17]. The observed decrease in lateral F-actin levels *in vitro* indicates that the intercellular junction defects under *Muc2* knockout are not reducible to secondary effects of inflammation or bacterial invasion: they persist in a culture devoid of immune cells and microbiota. Notably, the barrier function of organoids derived from *Muc2*^−^/^−^ animals remained intact in the functional permeability assay (Fig. 2a). Apparently, the alterations of lateral actin at this stage do not yet lead to barrier disruption and can be regarded as early molecular shifts preceding clinical manifestations. Under *in vivo* conditions, local weakening of the actin cytoskeleton reduces the resistance of the epithelium to mechanical stress (peristalsis, movement of contents), which, in the absence of the mucus layer, can lead to barrier damage and penetration of bacterial antigens.

Thus, chronic inflammation developing against the background of *Muc2* knockout leads to a persistent rearrangement of the intestinal epithelium: its spatial organization and cytoskeletal architecture change, and the stem cell pool is reduced. The organoid model derived from such animals represents a convenient platform for studying the mechanisms of barrier dysfunction in colitis outside the systemic influence of the organism. It allows the changes caused by the absence of the Muc2 protein in the epithelium itself to be separated from the contribution of the microbiota, immune cells, and systemic factors.

As a result of the comparative analysis of 3D organoids derived from the colons of *Muc2*^−^/^−^ and *Muc2*^+^/^+^ mice, it was established for the first time that *Muc2* knockout alters the morphofunctional properties of the intestinal epithelium independently of the microbiota and the immune system. Budding is suppressed in these organoids: they retain a spheroid shape throughout the entire culture period. This is accompanied by a twofold reduction of the Lgr5^+^ stem cell pool. The data obtained clarify the current understanding of the contribution of the epithelium to the pathogenesis of inflammatory bowel diseases and indicate the promise of the organoid model derived from *Muc2* knockout mice for preclinical testing of therapeutic approaches.

## COMPLIANCE WITH ETHICAL STANDARDS

All applicable international, national, and/or institutional principles of animal care and use were complied with. All procedures performed in studies involving animals were in accordance with the ethical standards approved by the legal acts of the Russian Federation and with the principles of the Basel Declaration. The experimental protocols involving laboratory animals were approved by the IMCB SB RAS Ethics Committee (protocol No. 02/21 of 04/08/2021).

This article does not contain any studies involving humans as research subjects.

## AUTHOR CONTRIBUTIONS

Concept and experimental design (J.V.P., E.N.K.), data collection (J.V.P., S.S.M., A.V.Z., E.N.K.), data processing (S.S.M., E.N.K.), manuscript writing and editing (J.V.P., S.S.M., A.V.Z., E.N.K.).

